# Structural Insights into Hearing Loss Genetics from Polarizable Protein Repacking

**DOI:** 10.1101/556258

**Authors:** M. R. Tollefson, J. M. Litman, G. Qi, R. J. Marini, C. E. O’Connell, M. J. Wipfler, H. V. Bernabe, W. T. A. Tollefson, T. L. Casavant, T. A. Braun, R. J. H. Smith, M. J. Schnieders

## Abstract

Hearing loss is associated with ~8100 mutations in 152 genes, and within the coding regions of these genes are over 60,000 missense variants. The majority of these variants are classified as ‘variants of uncertain significance’ to reflect our inability to ascribe a phenotypic effect to the observed amino acid change. A promising source of pathogenicity information are atomic resolution simulations, although input protein structures often contain defects due to limitations in experimental data and/or only distant homology to a template. Here we combine the polarizable AMOEBA force field, many-body optimization theory and GPU acceleration to repack all deafness-associated proteins and thereby improve average structure resolution from 2.2 Å to 1.0 Å based on assessment with MolProbity. We incorporate these data into the Deafness Variation Database to inform deafness pathogenicity prediction, and show that advanced polarizable force fields could now be used to repack the entire human proteome using the Force Field X software.

## Introduction

As the most common human sensory deficit, deafness impacts an estimated 360 million people globally (WHO data, http://www.who.int/pbd/deafness/estimates/en/index.html). Its cause is multifactorial, and with recent advances in the application of targeted genetic sequencing technology to clinical medicine, our understanding of genetic contributions to deafness has greatly advanced. The use of deafness-specific gene panels has changed the clinical paradigm in the evaluation of the deaf patient and is laying the foundation for personalized gene therapy to treat hearing loss.

The targeted genetic sequencing panel developed by our group, which we refer to as OtoSCOPE, includes 152 deafness-associated genes (1, 2). Its use enables us to identify an average of 545 variants per patient, which are curated in the publicly available deafness-specific database we purpose built called the Deafness Variation Database (DVD, Figure 1, Table S1 http://deafnessvariationdatabase.org) (3, 4). The DVD collates data from major public databases and uses criteria recommended by the American College of Medical Genetics and Genomics (ACMG) to classify every genetic variant as benign (B), likely benign (LB), variant of uncertain significance (VUS), likely pathogenic (LP) or pathogenic (P) based on collected evidence and curation by experts in genetic hearing loss. Of the ~800,000 variants in the genes included on OtoSCOPE that are listed in the DVD, more than 60,000 missense variations exist. Of these variants, ~4,000 are LP/P, ~38,000 are VUSs and ~18,000 are LB/B.

**Figure 1.**
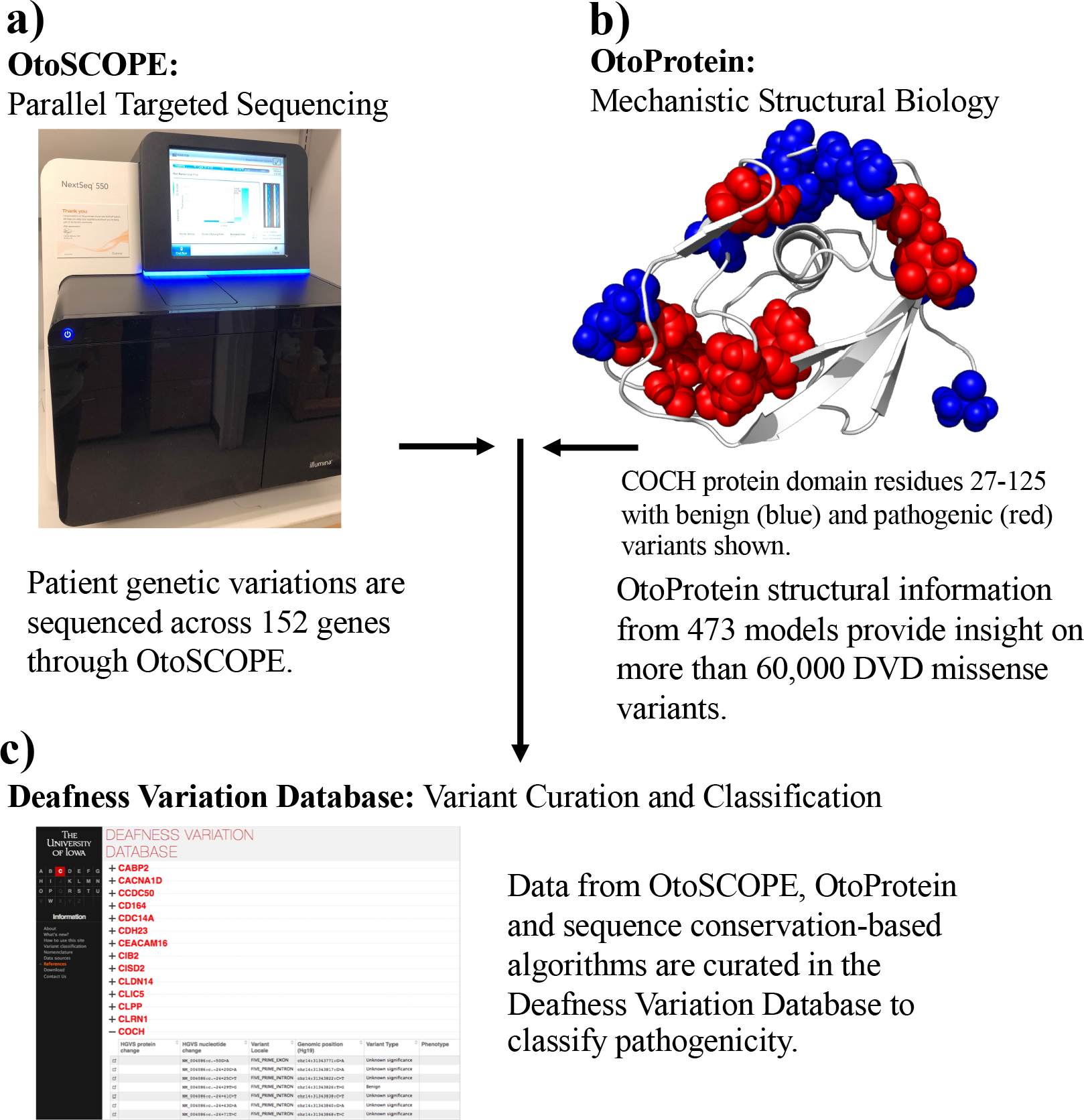
Incorporating Structural Insights into Variant Classification. a) OtoSCOPE sequencing technology discovers 545 variants per patient on average, 71 of which are non-synonymous coding, splice site variants or indel variants. b) Protein structural coverage for OtoScope genes is an important step toward identifying the molecular causation of disease-causing variants along with classifying VUSs. c) Variants collected through OtoSCOPE sequencing are curated in the Deafness Variation Database (DVD). The DVD combines minor allele frequency, experimental results, pathogenicity predictions from sequence conservation-based classifiers and now insights from protein structures (i.e. OtoProtein) for each variant. Nearly 80% of variants in the DVD remain unclassified and are assigned a pathogenicity of Variant of Uncertain Significance (VUS).

Many of the missense variations labeled as VUSs will ultimately be classified as LP/P, but we are currently relegated to classifying them as VUSs as a reflection of our inability to predict the phenotypic consequences of most genetic variations. We often lack variant-specific wet-lab-based functional evidence (5), and insights derived from simulations founded on protein structures must continue to mature to reliably make meaningful genotype-phenotype correlations.

Atomic resolution simulation techniques like molecular dynamics (MD) provide a promising first-principles approach for computationally predicting the potential impact of missense variants. However, its success is dependent on accurate protein structures. These structures are typically determined from an experimental method (*i.e.* X-ray crystallography, NMR, CryoEM, *etc*.) or from homology modeling. The latter leverages existing protein structure(s) as a template from which to create the model of a homologous amino acid sequence, and is most reliable when homologous sequences have at least 30% sequence identity as overall protein folding is typically conserved when this threshold is met (6, 7). To complement and enhance models available in databases such as ModBase (8) and SwissProt (9), dramatic improvements are possible by global optimization (*i.e.* repacking) of amino acid side-chains using more advanced molecular physics than was originally available (or could be computationally afforded) at the time of their creation.

For example, most protein structures found on both the Protein Databank (PDB) (10) and homology modeling databases (8, 9) are based on refinement with pairwise potential energy functions (*i.e.* force fields) such as the fixed charge Amber (11, 12), CHARMM (13, 14) and OPLS-AA (15, 16) models (17). Over the past decade, more accurate polarizable force fields have emerged that overcome limitations in previous generation pairwise models (18), including both the Atomic Multipole Optimized Energetics for Biomolecular Applications (AMOEBA) force field (19, 20) and the CHARMM Drude (21) model. Structural optimization with these state-of-the-art energy functions, when used with continuum representations of solvation (22–24), can compensate for limitations in experimental data and improve homology models. However, multiple challenges must be overcome to realize the benefits of polarizable force fields, including mitigating their increased computational expense and overcoming the loss of convenient pairwise approximations that are widespread in structural biology software such as Modeller (25), Phenix (26) and Rosetta (27).

Here we address these challenges to generate a family of deafness related protein structures called *OtoProtein*. Our approach combines the AMOEBA potential energy function (19, 20), many-body optimization theory (28), and GPU acceleration (29, 30) to optimize all available deafness-associated protein models. To assess the resulting structures objectively, we evaluated overall quality with the MolProbity (31, 32) algorithm. MolProbity identifies high-energy atomic clashes, unfavorable side-chain conformations and polypeptide backbone conformations inconsistent with low-energy secondary structure. The algorithm is widely used by crystallographers to aid refinement of models by reporting structural features that are known to be unphysical. Lower MolProbity scores are consistent with higher quality X-ray diffraction data (*i.e.* a score of 1.0 is calibrated to reflect 1.0 Å resolution data). Correcting rotamer outliers often improves other metrics and permits further relaxation of the structure with local minimization, resulting in more realistic, lower-energy structures for downstream analysis (*e.g.* molecular dynamics, alchemical free energy simulations, or feature extraction for bioinformatics analysis).

As described in the Results, our post-optimization OtoProtein dataset is near atomic resolution. These high-quality structures have been integrated with the DVD and are being used to define the structural impacts of deafness-related genetic variations as an aid to predicting variant effect and pathogenicity. Our polarizable protein repacking algorithm is freely available in the open source software Force Field X (FFX, http://ffx.biochem.uiowa.edu), and may be useful to others in the community that are integrating structural biophysics into variant classification.

## Methods

### A. Many-Body Energy Expansion Parallelization Across GPUs

Under a many-body potential, the total energy of a protein 𝐸(𝐫) can be defined to arbitrary precision using the expansion

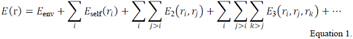

where *E*_env_ is the energy of the environment (*i.e.* the protein backbone and residues that are not being optimized), *E*_self_(*r*_*i*_) is the self-energy of residue *i* including its intra-molecular bonded energy terms and non-bonded interactions with the backbone, *E*_2_(*r*_*i*_, *r*_*j*_) is the 2-body non-bonded interaction energy between residues *i* and *j* with other residues turned off, and *E*_3_(*r*_*i*_, *r*_*j*_, *r*_*k*_. is the 3-body non-bonded interaction energy between residues *i*, *j*, and *k* with other residues turned off. The self, 2-body, and 3-body energy terms are calculated as follows, where *E*_*BB*/*SC*_ is the total energy of the backbone with the sidechain(s) of the selected residue(s) included (shown graphically in Figure 2a).

**Figure 2.**
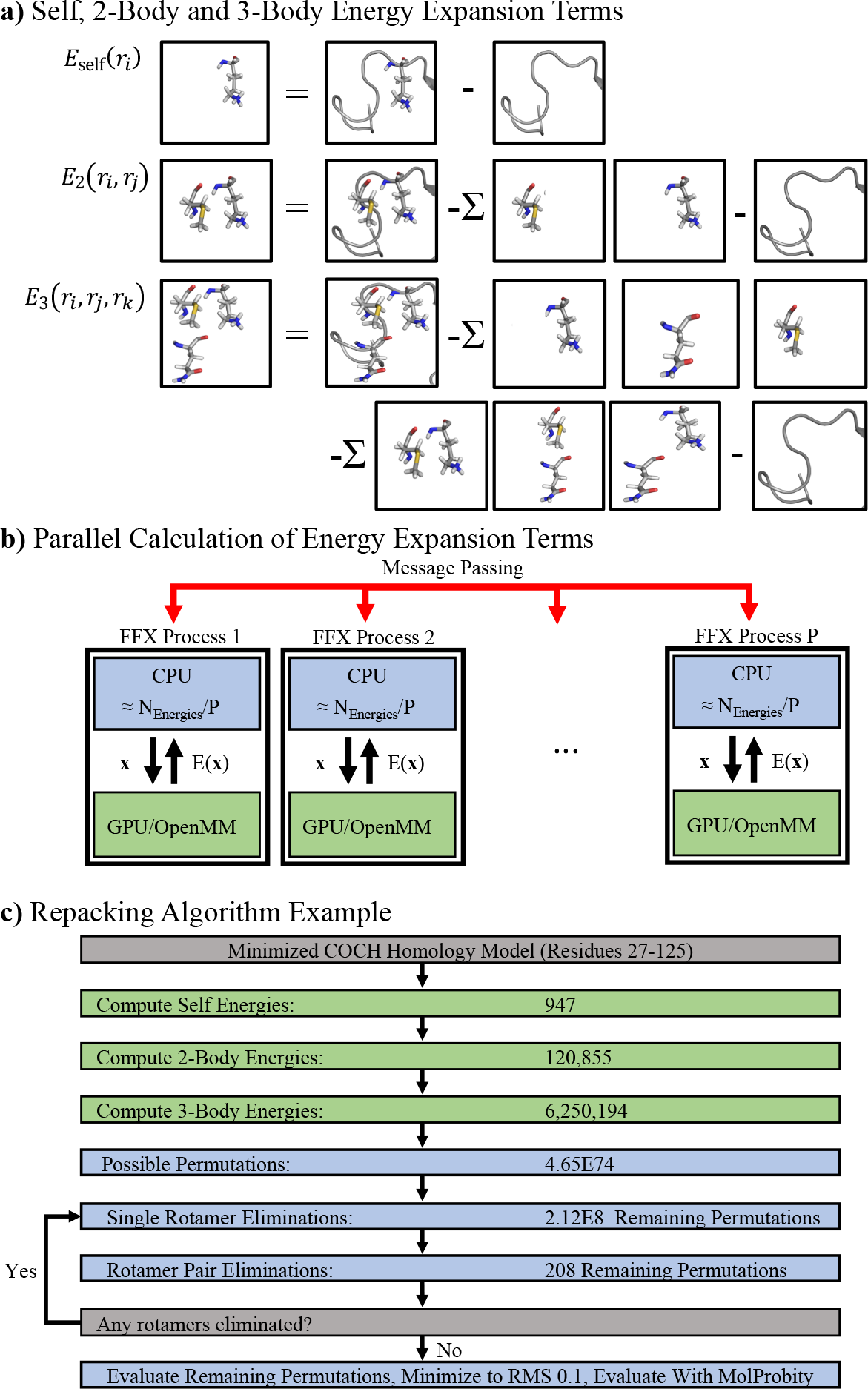
Overview of the protein repacking algorithm. a) Depiction of rotamer self, 2-body and 3-body energy terms. b) Parallel computation of energy terms across processes and GPUs. Processes (blue boxes) are each assigned a group of self-energies to calculate, where 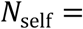 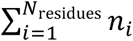 is the sum of rotamers across all residues to give *N*_self_/*P* evaluations per process. Processes compute energy values by sending a conformation **x**to a GPU (green box) for evaluation using the OpenMM API, followed by return of the energy E(**x**) and its communication to all processes using Java MPI (red arrows). The 2-body and 3-body energies are parallelized in a similar fashion. c) The number of side-chain energies and conformational permutations for a 98 residue COCH protein domain are shown as an example. After all energy terms have been calculated (green rectangles), the combinatorial side-chain conformational space is reduced using many-body Goldstein rotamer and rotamer pair elimination criteria (see Methods) to achieve a tractable number of permutations to evaluate. Prior to eliminations, 4.65*1074 side-chain permutations exist, but only 208 permutations remain to be evaluated after eliminations.

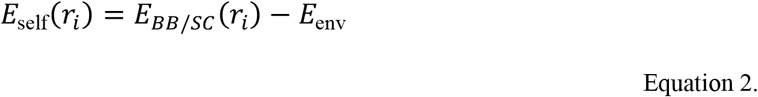

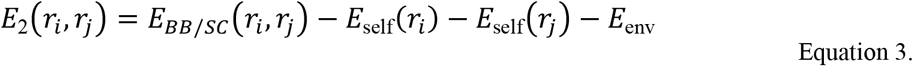

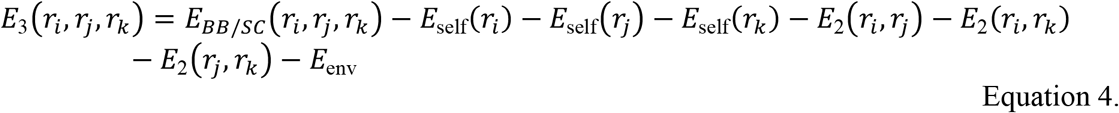

Individual energy evaluations are calculated on graphical processing units (GPUs) *via* the CUDA kernels of OpenMM (30), and the evaluations are distributed over many GPUs, potentially over multiple nodes, using the Parallel Java library (33) (Figure 2b). Side-chain rotamer conformations that are not part of the optimum structure can be rigorously eliminated using mathematical expressions (28, 34, 35) (Figure 2c).

Computing the self, 2-body and 3-body energy terms as a function of rotamer conformation is computationally expensive. To address this challenge, our Force Field X (FFX) program (36) utilizes two complementary parallelization approaches, including 1) use of the Parallel Java (33) (PJ) message passing interface (MPI) library to distribute terms among multiple processes, and 2) use of the OpenMM API (30) to perform force field energy evaluations on NVIDIA GPUs *via* CUDA kernels. FFX uses PJ to divide each shared memory node of a multiple node compute cluster into one or more processes (Figure 2b). Energy terms are then assigned to processes, evaluated and globally communicated across all processes using PJ message passing, with synchronization steps between calculation of the self, 2-body, and 3-body energy terms (*i.e.* 2-body terms depend on self-terms as shown in Equation 3, and thus must be calculated after self-energies are completed and before 3-body energies). The FFX-OpenMM interface (based on Java Native Access wrappers to the OpenMM C++ API) can be used to offload energy evaluations from FFX, which executes on CPUs, to OpenMM on a GPU. Once all energy terms are calculated, side-chain rotamers and rotamer pairs are eliminated by lower energy alternatives based on rigorous mathematical inequalities that have been described for pairwise force fields (*e.g.* dead-end elimination (34) and Goldstein elimination (28, 35)) and more recently generalized to include 3-body terms for use with many-body force fields (28) such as the polarizable atomic multipole AMOEBA model (20, 37). The many-body Goldstein criteria for rotamer elimination (28), truncated at 3-body terms, is given by

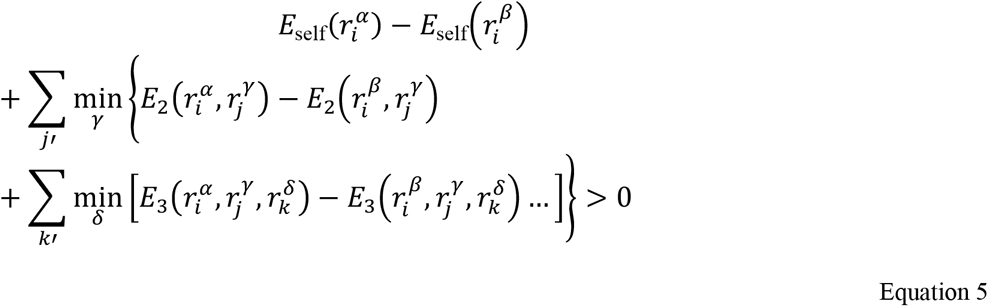

and if satisfied indicates rotamer *α* of residue *i* is eliminated by rotamer *β* (the ellipses signify the presence of further higher-order terms). The expression for rotamer pair elimination is given by

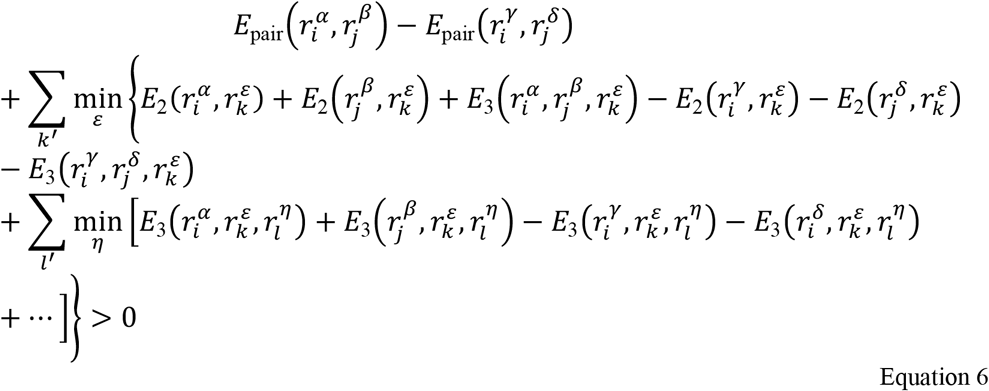

and if satisfied indicates that the rotamer pair 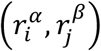 for residues *i* and *j* is eliminated by rotamer 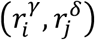.

Four approximations to rigorous use of the many-body Goldstein inequalities given above were explored, each of which is summarized here and described more fully in the Results section. First, it was determined that the expansion could be truncated at pairwise terms, due to damping of 3-body and higher order terms by the generalized Kirkwood implicit solvent. However, in the absence of implicit solvent, previous work demonstrated that inclusion of 3-body terms is sometimes necessary (28). The second approximation was a distance cutoff; if the closest rotamers for a residue pair or triple are more than 2 Å apart, the interaction energy is set to 0. Third, pruning was utilized to remove rotamers with self-energies 25 kcal/mol or more above the lowest self-energy of a residue, before calculation of 2-body energies. This criterion is based on the heuristic observation that rotamers with such an unfavorable self-energy *(e.g.* due to an atomic clash with backbone atoms) are unlikely to be part of a well-packed structure. However, for structures with significant backbone flaws, this approximation must be used with care because it can incorrectly eliminate the "least bad" rotamer that is actually part of the global minimum conformation. Our final approximation involved imposing a 3D grid over the protein, followed by optimization within each cube, rather than including all protein residues simultaneously. Although the repacking algorithm is a provable global optimizer within a single cube of the grid, it is not for the protein grid as a whole because coordinated changes between cubes are neglected.

### B. The *OtoProtein* Structure Database

Comparative protein modeling provides a means to predict the structure of a protein whose atomic coordinates have not been solved experimentally by crystallography, NMR, *et cetera* (38). Many human genes implicated in hearing loss have not been studied experimentally, so computational approaches are necessary to generate high quality protein structures. Comparative protein modeling begins from an experimental structure for an evolutionarily related protein, which is used as a template for the target sequence (10, 39). The percent sequence identity between the homologues provides an estimate of model reliability (40). Comparative protein models are conducive to the study of protein function, dynamics, and interactions with other molecules such as ligands, DNA, RNA, or other proteins. Homology models can also be used to study missense variants, providing a promising basis for understanding the role of protein phenotypes in heterogenic diseases like hearing loss.

However, comparative protein models from leading databases often include defects directly related to approximations in the methods used for their generation (*e.g.* pairwise force fields, local rather than global optimization, *etc*). We sought to improve comparative protein models from SwissProt (39) and ModBase (8) spanning 152 genes included in the OtoSCOPE platform. Although using homology models based on a sequence identity of 30% or greater generally gives confidence the protein backbone fold has been evolutionarily conserved (40), this work includes all publicly available models (the average sequence identity was 41.7% for all 472 structural models). Both SwissProt and ModBase strive to provide structural coverage for the largest portion of the human proteome possible, however, this limits their ability to explore the use of advanced many-body force fields. Here we show that use of the polarizable AMOEBA force field in tandem with global optimization of amino acid side-chains (28) can significantly improve the quality of SwissProt or ModBase structures as assessed by tools like MolProbity (31, 32). High quality protein structural models, in turn, provide optimal starting points for downstream molecular dynamics algorithms that can be used to analyze missense variations. The parallelized repacking algorithm described here demonstrates that it is now feasible to refine large databases of homology models using advanced polarizable force fields.

All homology models were refined using the 2018 AMOEBA force field (20, 41) with Generalized Kirkwood implicit solvent (23). The input homology models were first locally optimized using the L-BFGS algorithm available in FFX that is accelerated using OpenMM on GPUs to an RMS gradient convergence criterion of 0.8 kcal/mol/Å. The rationale for minimizing with a relatively loose convergence criterion prior to rotamer optimization was to relax the backbone conformation without excessively favoring the starting conformation over alternative rotamers. Locally optimizing to a tighter convergence criterion prior to side-chain optimization resulted in higher energy, less favorable structures due to over-stabilizing starting rotamers. Next, the side-chain repacking algorithm was applied, followed by a final local L-BFGS minimization to an RMS gradient convergence criterion of 0.1 kcal/mol/Å. The resulting protein structures and original homology models were then evaluated and compared using both the MolProbity assessment tool and AMOEBA/GK energies.

## Results

### A. Polarizable Protein Repacking Algorithm Using GPUs

To benefit fully from the emergence of polarizable force fields in the context of protein structure prediction and repacking, the theory that underlies established algorithms must be revisited in order to incorporate many-body electronic polarization and to optimize performance across GPUs. We examined four approximations to the polarizable protein repacking algorithm to enhance efficiency, while maintaining structure quality. The approximations are illustrated using a 98-residue homology model of the COCH protein (residues 27-125), which is based on an NMR template with 98% sequence identity. Previous work showed that truncating the energy expansion at 3-body terms resulted in accurate side-chain positions being identified in the context of real space X-ray refinement (28). However, when using the Native Environment Approximation (NEA) (42) in combination with the AMOEBA generalized Kirkwood implicit solvent (23), we found that the contribution of energy terms within the energy expansion decays quickly and that truncation at 2-body terms is sufficiently accurate for repacking in implicit solvent (Table S2). Rapid decay is due to implicit solvent damping 3-body electrostatics to such an extent that they generally do not affect side-chain rotamer eliminations (whereas our prior work did not employ an implicit solvation model). Truncation at 2-body energy terms results in a nearly 52x speed-up (Table S3) as compared to the original rotamer optimization protocol (28) without any rotamer changes compared to including 3-body terms (Table 1a). In future work, we plan to additionally optimize the protonation states of all titratable residues, which will necessitate a fresh appraisal of the impact of 3-body energies due to the formal charge of residues changing.

**Table 1.** Adjustable repacking parameters are examined in the context of computational expense and structural quality. All tests used residues 27-125 of isoform 1 of the COCH protein. All relative potential energies (in kcal/mol) are compared to the global minimum of the COCH protein as calculated when using no approximations. Times are wall clock times in seconds using a node with 4 GPUs. For each adjustable parameter, the recommended choice for use with AMOEBA and the GK implicit solvent is shown in bold. The individual and overall speed-ups for each approximation are given.

| **A) Truncation of the energy expansion at either 2-body or 3-body terms.** |  |  |  |  |
| --- | --- | --- | --- | --- |
| Expansion Truncation | Relative Energy | Time | Speed-Up |  |
| **2-Body** | 0.0 | 847 | 51.7x |  |
| 3-Body | 0.0 | 43850 | 1.0x |  |
| **B) Based on truncation at 2-body terms, distance cutoffs from 1 to 6 Å are evaluated. Criteria for evaluating cutoffs are described in the main text.** |  |  |  |  |
| Distance Cutoff (Å) | Relative Energy | Time | Speed-Up | Overall |
| 1 | 32.6 | 137 | 5.9x | 320.0x |
| **2** | 0.2 | 304 | 2.7x | 144.2x |
| 3 | 0.0 | 420 | 1.9x | 104.4x |
| 6 | 0.0 | 813 | 1.0x | 53.9x |
| **C) Based on truncation at 2-body terms and a 2 Å distance cutoff, pruning thresholds are evaluated.** |  |  |  |  |
| Pruning Threshold | Relative Energy | Time | Speed-Up | Overall |
| 5 | 0.2 | 43 | 7.1x | 1019.8x |
| 15 | 0.2 | 113 | 2.7x | 388.1x |
| **25** | 0.2 | 142 | 2.1x | 308.8x |
| No pruning | 0.0 | 304 | 1.0x | 144.2x |
| **D) Based on truncation at 2-body terms, a 2 Å distance cutoff, and 25 kcal/mol pruning threshold, cube edge lengths from 10 to 30 Å are evaluated for cube optimization.** |  |  |  |  |
| Cube Size (Å) | Relative Energy | Time | Speed-Up | Overall |
| **10** | 0.2 | 40 | 3.6x | 1096.3x |
| 20 | 0.9 | 94 | 1.5x | 466.5x |
| 30 | 0.2 | 142 | 1.0x | 308.8x |

The second approximation applies a distance-based cutoff between residues, which results in the interaction energy of two or more side-chains being set to 0 if the minimum atomic distance between rotamer permutations is above a defined cutoff. The minimum distance *d*min between two residues *i* and *j* is calculated using the expression

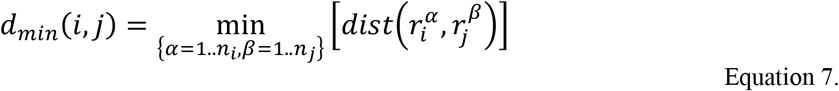

where the min operation is over the set of all rotamer permutations (*i.e.* residues *i* and *j* have *n*_*i*_ and *n*_*j*_ rotamers, respectively, to give *n*_*i*_ × *n*_*j*_ permutations), and the distance function (*dist*) returns the minimum pairwise atomic distance given rotamer conformations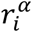 and 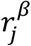. We tested a range of cutoffs and found that 2 Å, when combined with truncation at 2-body energies, provides a 144.2x speed-up compared to the original protocol while only increasing the energy relative to the global minimum by 0.2 kcal/mol (*i.e.* two solvent-exposed side-chains had different conformations) (Table 1b). Although 2 Å is an aggressive cutoff, evidence from our dataset of protein models (Table S4) suggests structures still closely approach the global minimum, but at much less expense.

The third approximation prunes rotamers and/or rotamer pairs if their conformation is higher than the lowest energy alternative plus a threshold

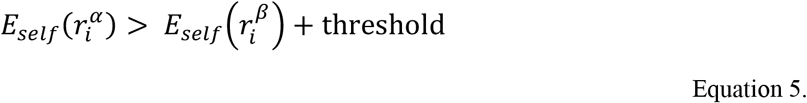

A pruning threshold of 25 kcal/mol results in further speed-up without compromising the quality of output structures (Table 1c). Although pruning inequalities are not rigorous, unlike the mathematically proven Goldstein eliminations (see Methods), they obviate calculating many pair energies to yield over a 3x speed-up. Pruning did not result in any additional changes to rotamer conformations as compared to the global minimum found when using a 2-body expansion and cutoff of 2 Å.

The final approximation uses a series of cube-shaped domains defined by imposing a 3D grid over the protein followed by sequential optimization of each cube of the grid. This approximation is especially useful for large protein domains that have an intractable number of energetically closely spaced permutations even after application of elimination criteria. By varying cube size and cube overlap, we determined that a cube edge length of 10 Å with no overlap optimized performance without degrading quality (Table 1d). Cube optimization results in no additional change in energy relative to the global minimum found when using a 2-body expansion and cutoff of 2 Å (note that a 30 Å cube contains the whole COCH domain and is a global optimization). Combining all four optimal approximations results in a total speed-up of approximately 3 orders of magnitude.

We next implemented a parallelization approach that combines Parallel Java with GPU acceleration. As the number of nodes is increased, our Parallel Java message-passing parallelization algorithm achieved a near linear speed-up (Tables 2 and S5). Offloading energy evaluations to OpenMM on a single node equipped with a GPU (two Intel Xeon E5-2680v4 CPUs and one NVIDIA GTX 1080 TI GPU) resulted in a 11.5-fold speed-up compared to using the same node with no GPU (*i.e.* a single GPU was 11.5x faster than parallelization over all 28 Intel CPU cores). Our original CPU parallelized Java implementation of the algorithm with no approximations (same protein domain run on two Intel Xeon E5-2680v4 CPUs, 3-body expansion, 6Å cutoff, and no pruning) required calculation of over 6 million energy evaluations and consumed 16.5 compute days on a node. By combining algorithm approximations, parallelization across 4 processes on one node (*i.e.* PJ message passing) and GPU acceleration (1 GPU per process, 4 GPUs total), our algorithm executes the 20,232 necessary energy evaluations in 142 seconds.

**Table 2.** Energy evaluation timings for global side-chain optimization of COCH residues 27-125 using a varying number of GPUs (each node contains two Intel Xeon E5-2680v4 CPUs and 4 nVidia GTX 1080 TI GPUs).

| Number of Nodes | Number of GPUs | Time for Energies (sec) | Speed-Up (Relative to Using all CPU Cores) |
| --- | --- | --- | --- |
| 1 | 0 (CPUs only) | 5505 | 1.0x |
| 1 | 1 | 479 | 11.5x |
| 1 | 2 | 251 | 21.9x |
| 1 | 4 | 142 | 38.8x |
| 2 | 8 | 76 | 72.4x |
| 4 | 16 | 43 | 128.0x |

### B. The*OtoProtein* Structure Database

We applied our accelerated repacking algorithm to a set of 473 deafness-associated protein models. For both starting homology models and refined structures, quality was assessed using the heuristic MolProbity algorithm that examines steric clashes, poor side-chain rotamers and amino acid backbone favorability (*i.e.* phi/psi dihedral angle combinations). The MolProbity score is calibrated to predict the quality of X-ray diffraction data that is expected to have produced the assessed structure (*i.e.* a MolProbity score of 1.5 corresponds to an expected X-ray resolution of 1.5 Å, where lower values indicate higher quality). On average, we reduced steric clashes per 1000 atoms from 25.1 to 0.03, decreased Ramachandran outliers from 2.03% to 0.94%, and decreased the percentage of poor side-chain rotamers from 2.3% to 1.6% (Table 3). Overall, the repacking protocol improved the mean MolProbity score from 2.16 Å to 1.04 Å, consistent with protein structural models at atomic resolution (Figure 3). The average AMOEBA force field energy for the dataset when locally optimized to RMS gradient convergence criteria of 0.8 kcal/mol/Å, was −15,342 kcal/mol. After global side-chain optimization, average AMOEBA energy for the dataset was reduced to −16,287 kcal/mol, a reduction of 945 kcal/mol from the structures that were minimized to an RMS gradient criterion of 0.8 kcal/mol/Å without rotamer optimization. Although local minimization without rotamer optimization dramatically reduces atomic clashes, the number of poor rotamers increased from 2.3% to 2.9% and motivates the need for side-chain repacking. The overall repacking procedure required just 71 GPU-days for all 473 OtoProtein structures. The complete list of statistics for each model is available in Table S6. Based on these results, GPU-accelerated repacking with the polarizable AMOEBA force field could potentially be used to improve the quality of large protein structure databases with only a modest investment in hardware.

**Figure 3.**
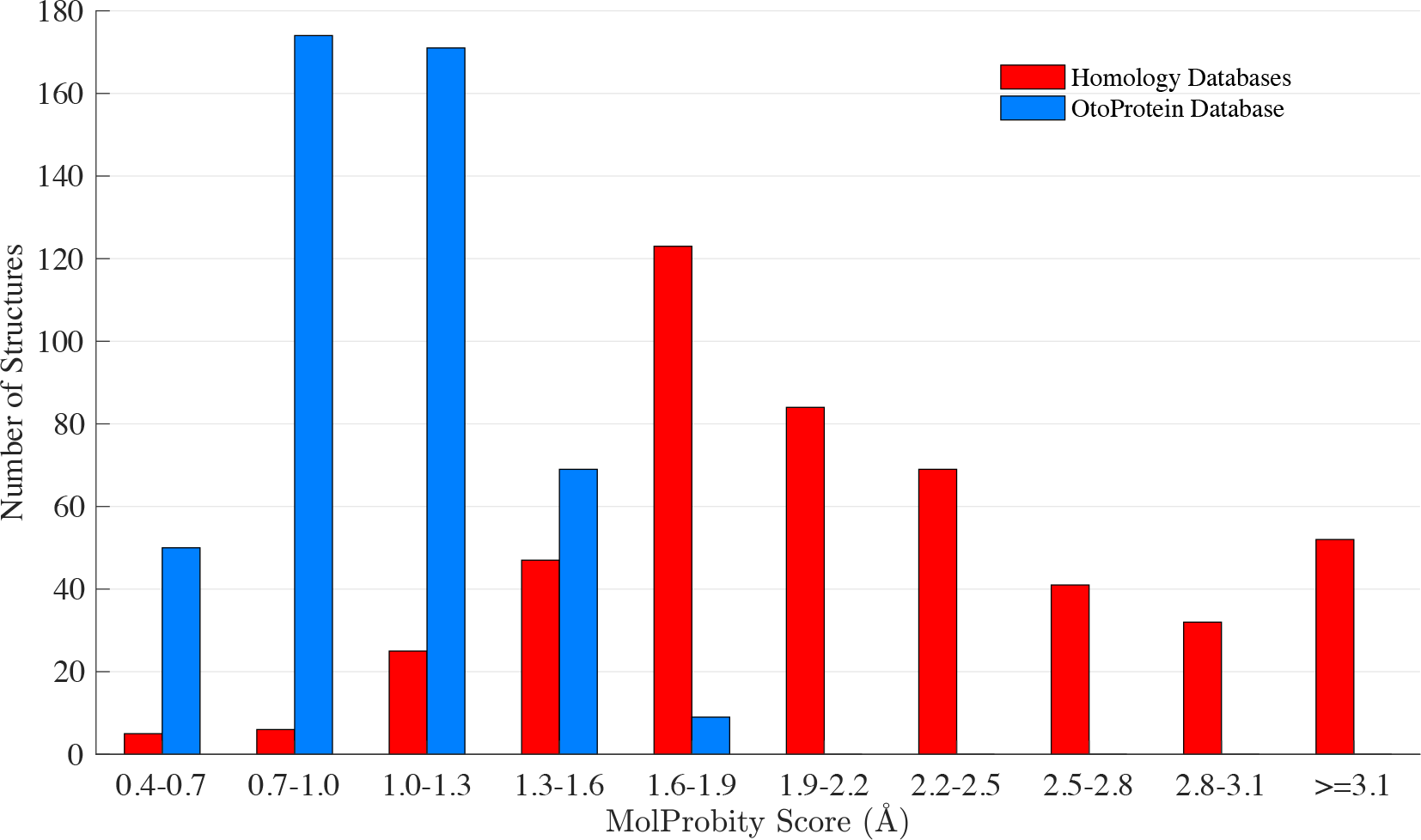
Histogram of MolProbity resolutions for the OtoProtein Structure Database before and after optimization. Before optimization (red), the 473 structures have an average MolProbity score of 2.16 Å, while after optimization (blue) the dataset has an average MolProbity score of 1.04 Å (*i.e.* approaching the quality expected of atomic resolution X-ray structures).

**Table 3.**
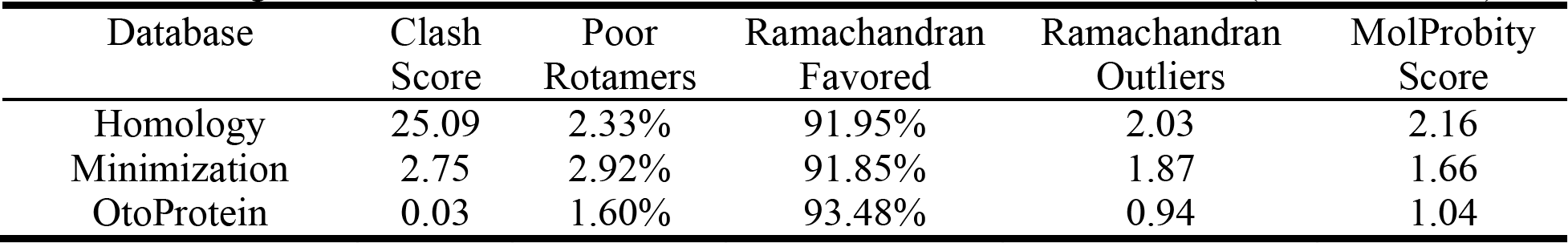
Average refinement statistics for the OtoProtein Structure Database (473 structures).

The OtoProtein Structure Database has been incorporated into the DVD to provide public availability of the models in combination with the exhaustive DVD genetic information. The combination of OtoProtein structural information with existing DVD data (*e.g.* minor allele frequency, pathogenicity assessment, *etc*.) provides a powerful platform for the auditory research community. For example, it is now possible to visualize and gain insight on the clustering of pathogenic variations in specific domains of a protein and to examine structural features that correlate with pathogenicity (Figure 4).

**Figure 4.**
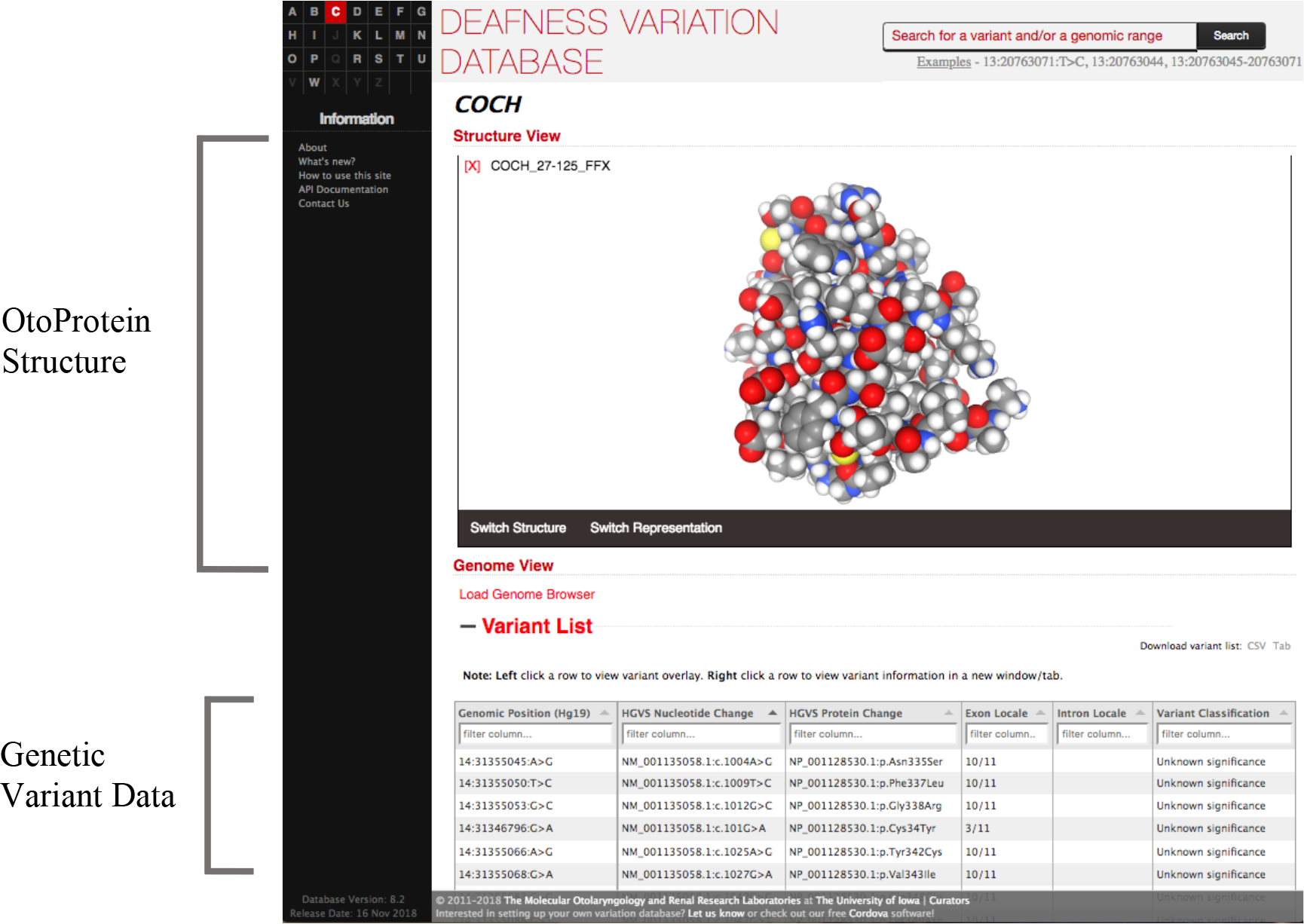
Incorporation of OtoProtein into the Deafness Variation Database (DVD). All models developed with the GPU-accelerated polarizable protein repacking algorithm are publicly available in the DVD, where they can be viewed in combination with genomic and variant data.

## Discussion

Structural coverage of the human proteome has increased rapidly since the early 1990s, with ~40% of the human proteome now having comparative models based on templates with a sequence identity of at least 30% (9). Here we applied a GPU accelerated polarizable protein repacking algorithm to the deafness-associated proteome defined by homology models of any sequence identity (average sequence identity of the dataset is 41.7%). We found that 38.8% of the deafness-associated proteome could be modeled structurally, comparable to structural coverage of the entire human proteome. The 473 structural models we collected and optimized span 145 deafness-associated genes. These structures had an initial average MolProbity score of 2.16 Å, but after repacking, the average score improved to approximately atomic resolution at 1.04 Å. These calculations required just 71 GPU-days. In addition to covering nearly 40% of OtoSCOPE with atomic resolution structural models, our OtoProtein database provides structural coverage for 22,809 of the 61,971 missense variations in the DVD (16,203 are VUSs, 1,931 are LP/P, and 4,675 are B/LB). These models are publicly available in the DVD through the NGL protein viewer (43). When integrated with information on patient missense variations available through the DVD, the OtoProtein database represents a unique tool for understanding deafness genetics from a structural perspective. Building on the OtoProtein structural platform, future work will model missense variations using thermodynamic free energy simulations to provide insights on how variants disrupt protein folding and/or protein-protein interactions.

The GPU-accelerated protein repacking algorithm is freely available to the research community through the Force Field X program, which may be useful to refine other structural datasets outside of the deafness domain. The algorithm is designed for use with advanced polarizable force fields and features an energy expansion up to 3-body interactions. Computational speed is achieved using an architecture based on parallelization across an arbitrary number of compute nodes and GPUs, and together with repacking algorithm optimizations provides multiple orders of magnitude speed-up without compromising structural quality. Although polarizable repacking algorithms were previously not efficient enough to apply to large-scale datasets, this work opens the door to their application to all models of the human proteome. The Swiss Model Repository (SMR), for example, lists 45,083 homology models with an average residue length of 232 amino acids (9). Structures of this size (*i.e.* ~230 residues) require only ~260 seconds to repack using our algorithm on a node with 4 GPUs (*e.g.* repacking our DSPP 88-318 model of 230 residues took 262 seconds of wall clock time). Based on the average model size in the SMR, we estimate that repacking all SMR human proteins would require only ~140 days on a node equipped with 4 GPUs (*i.e.* ~2 weeks on our compute cluster that has 10 such nodes).

Finally, a limitation to the current repacking algorithm is its reliance on existing homology models to serve as initial coordinates. While it can greatly improve the quality of existing structural models, it does not provide coverage of proteins through *ab initio* or *de novo* techniques. This limitation is the subject of on-going work based on GPU-accelerated biased sampling methods, which we are using to expand structural coverage of the OtoSCOPE proteome. Despite this limitation, the OtoProtein structural information is already being used to gain insight into the protein phenotype of more than 20,000 missense variants associated with deafness.

## Author Contributions

Conceived the theory: MRT, JML, MJS. Performed the experiments: MRT, JML, GQ, CEO, WTAT. Analyzed the data: MRT, JML, RJHS, MJS. Contributed code/tools/structures: MRT, JML, GQ, CEO, RJM, HVB, WTAT, TAB, TLC, MJS. Wrote the paper: MRT, JML, WTAT, TAB, TLC, RJHS, MJS.

## Acknowledgements

All computations were performed on The University of Iowa Argon cluster with support and guidance from Glenn Johnson, Ben Rogers and Brenna Miller. This material is based upon work supported by the National Science Foundation Graduate Research Fellowship under Grant No. 000390183. Any opinion, findings, and conclusions or recommendations expressed in this material are those of the authors(s) and do not necessarily reflect the views of the National Science Foundation. JML was supported by NIH/CBB Award T32 GM0008365 and the University of Iowa Presidential Graduate Research Fellowship. MJS was supported by NIH RO1DK110023, NIH RO1DC012049, and NSF CHE-1751688. GQ and CEO were supported by fellowships from The University of Iowa Center for Research by Undergraduates.

